# Effects of Global *Ripk2* Genetic Deficiency in Aged Mice following Experimental Ischemic Stroke

**DOI:** 10.1101/2025.02.05.636687

**Authors:** John Aaron Howell, Jonathan Larochelle, Rachel E. Gunraj, Sofia M. Stansbury, Lei Liu, Changjun Yang, Eduardo Candelario-Jalil

## Abstract

Besides the loss of blood and oxygen reaching the ischemic tissue, many secondary effects of ischemic stroke can cause additional tissue death, including inflammation, oxidative stress, and proteomic disturbances. Receptor-interacting serine/threonine kinase 2 (RIPK2) is an important mediator in the post-stroke inflammatory cascade that responds to signals and molecular patterns released by dead or dying cells in the ischemic area. We hypothesize that RIPK2 signaling worsens injury and neurological recovery post-stroke and that global deletion of *Ripk2* will be protective following ischemic stroke in aged mice. Aged (18-24 months) male mice were subjected to permanent middle cerebral artery occlusion (pMCAO). Vertical grid, weight grip, and open field were conducted at baseline and on days 1, 2, 3, 8, 15, and 22 post-stroke. Cognitive tests (novel object recognition and Y-maze) were performed at baseline and day 28 post-stroke. Infarct size was measured using cresyl violet staining, and reactive gliosis was measured using Iba1 and GFAP staining at day 28 post-stroke. Global deletion of *Ripk2* (*Ripk2^-/-^*) in aged mice resulted in smaller infarct volume and improved performance on vertical grid and weight grip tests compared to aged wildtype (WT) mice. Additionally, aged *Ripk2*^-/-^ mice had less Iba1 staining in the ipsilateral cortex than the aged WT control mice. This study further elucidates the role of RIPK2 signaling in the ischemic cascade and expands our knowledge of RIPK2 in stroke to aged mice. These results support the hypothesis that RIPK2 signaling worsens injury post-stroke and may be an attractive candidate for therapeutic intervention.

## Introduction

Stroke is the fifth leading cause of death in the United States and a major cause of long-term disability. Ischemic strokes, caused by a blood clot in a major cerebral artery, comprise 87% of total stroke incidence [1]. Currently, therapeutic interventions, including recombinant tissue plasminogen activator (rtPA), Tenecteplase (TNKase), and mechanical thrombectomy, only aim to remove the clot to allow for revascularization of the ischemic tissue [2,3]; however, there are many secondary effects of ischemic stroke that can lead to additional tissue death and worsened neurological outcomes long-term. In addition to the lack of oxygen and energy reaching the tissue, ischemic strokes cause increased inflammatory signaling, oxidative stress, and excitotoxicity that worsen the injury and are not ameliorated by currently approved treatment options [4]. Thus, there is a great need for novel therapeutics in stroke to target these secondary mechanisms to reduce injury and improve long-term recovery post-stroke.

Neuroinflammation starts moments after ischemia begins [4,5]. As neurons die within the ischemic core, microglia and astrocytes are activated, leading to the release of pro-inflammatory cytokines and the formation of reactive oxygen species and excitotoxicity [6]. The release of cytokines and chemokines weakens the blood-brain barrier (BBB), which results in infiltration of peripheral immune cells that can exacerbate tissue injury [4]. Neuroinflammation can last for days to weeks following a stroke and can cause additional tissue injury and worsened neurological outcomes [7,8].

Receptor-interacting serine/threonine kinase 2 (RIPK2) is an important mediator in inflammatory signaling [9,10]. Following injury or disease, cells release pathogen-associated molecular patterns (PAMPs) or damage-associated molecular patterns (DAMPs), which are recognized by nucleotide-binding oligomerization domains (NOD) 1 and NOD2 receptors. When NOD1/2 recognize PAMPs or DAMPs, they become activated and oligomerize to recruit RIPK2, which is subsequently ubiquitinated. RIPK2 then can activate transforming growth factor-β (TGF-β)-activated kinase 1 (TAK1) and inhibitor of κB kinase (IKK), leading to nuclear factor-kappa B (NF-κB) and mitogen-activated protein kinases (MAPK) activation and inflammatory signaling [9,10]. RIPK2, being upstream of multiple inflammatory cascades, is an attractive candidate for pharmacological therapies targeting inflammation.

RIPK2 has been shown to be involved in the progressive neuroinflammation associated with multiple sclerosis and intracerebral hemorrhage [11,12]. RIPK2 has also been implicated in many inflammatory gastrointestinal disease states, such as Crohn’s disease, ulcerative colitis, and inflammatory bowel disease [13,14]. Previous work in our lab has elucidated a clear role of RIPK2 signaling in the neuroinflammatory response following rodent models of ischemic stroke. Using pharmacological inhibitors, we have shown that inhibiting RIPK2 *in vivo* results in reduced infarct volumes and improved behavioral outcomes in young mice [15]. We have also shown that both global knockout and microglia-specific conditional knockout of *Ripk2* results in smaller infarct volumes and improved behavioral outcomes, implicating the microglia as a primary driver of the detrimental effects of RIPK2 signaling post-stroke [16]. However, these studies were done in younger animals, and since stroke risk increases with aging [1], it was important to continue this work using aged animals.

Stroke prevalence and the likelihood of death due to stroke-related complications increase with age [1]. Additionally, aging leads to heightened neuroinflammation through cellular senescence and changes in immune signaling [17]. Because of these factors, advanced age is a critical biological variable to include in animal models of ischemic stroke. This study aims to expand the data showing the role of RIPK2 signaling in stroke by using aged global *Ripk2* knockout mice and assessing long-term motor and cognitive behavior, as well as infarct size and gliosis measurements. We hypothesized that the aged *Ripk2* deficient mice would have smaller infarction volumes and improved behavioral outcomes following permanent middle cerebral artery occlusion (pMCAO), a rodent model of ischemic stroke.

## Materials and Methods

### Animals

Mice were housed in a specific pathogen-free facility with a 12-hour light/12-hour dark cycle and ad libitum access to food and water. Mice deficient for the *Ripk2* allele (Jackson Laboratory, stock #007017) were aged in-house to 18-24 months of age. Wildtype (WT) control aged animals (18 mo) were obtained from the National Institute of Aging (NIA) and acclimated to our facility for several weeks before the experiments. All animal experiment procedures were conducted following the NIH Guide for the Care and Use of Laboratory Animals. All procedures were approved by the University of Florida Institutional Animal Care and Use Committee (animal protocol numbers 202200000201). All experiments and analyses were performed by investigators blinded to animal genotypes. The specific number of animals for each analysis is stated in the figure legends.

### Permanent Middle Cerebral Artery Occlusion (pMCAO)

Permanent cerebral ischemia was induced by ligation of the left middle cerebral artery (MCA), as described previously [18]. In short, mice were anesthetized with 1.5-2 % isoflurane in medical-grade oxygen. The left common carotid artery (CCA) was exposed and carefully dissected from the vagus nerve and a 7-0 silk suture was applied to ligate the left CCA permanently. A skin incision was made between the eye and the left ear under a stereomicroscope, and the temporal muscle was retracted to locate the MCA via skull transparency. A small round craniotomy (∼1-1.5 mm in diameter) was made between the zygomatic arch and the squamosal bone to expose the MCA using a surgical burr (Cat No. 726066) connected to a battery-operated Ideal Micro-Drill (Cat No. 726065; Cellpoint Scientific, Gaithersburg, MA). Sterile saline was applied to the target area, and the meninges covering the MCA were carefully removed using forceps. The MCA distal trunk was permanently ligated using an ophthalmic 9-0 silk suture just before the bifurcation between the frontal and parietal branches of the MCA. A clear interruption of blood flow was visually observed. After surgery, mice were allowed to recover in a temperature-controlled chamber.

### Cresyl Violet Staining and Immunohistochemistry

Mice were euthanized at 28 days post-stroke and transcardially perfused with 10 mL of saline containing 5 mM EDTA using a peristaltic pump at a speed of 5 mL/min, followed by 30 mL of 4 % PFA in PBS. Brains were collected and post-fixed in 4 % PFA for 24 hours, then transferred to 30 % sucrose at 4°C until they sank. Brains were cut coronally into a series of 30-μm thick sections using a cryostat.

### Cresyl Violet Staining

Cresyl violet staining was performed to measure infarct volume at 28 days post-pMCAO, as described in our previous work [19]. Sections were scanned using an Aperio ScanScope® CS system and analyzed with ImageScope Software (Aperio Technologies; Vista, CA). The border between infarcted (dark purple stain) and non-infarcted area (light purple stain) was outlined and quantified using the software. Infarct volume was calculated by subtracting the area of healthy tissue on the ipsilateral hemisphere from the total area of the contralateral hemisphere.

### Immunohistochemistry

IHC was performed as previously described [16,20]. Sections were stained with either rabbit anti-Iba1 (1:5000; catalog # 019-19741, Wako Bioproducts; Richmond, VA) or rabbit polyclonal glial fibrillary acid protein (GFAP; 1:3000; catalog # Z0334, DAKO, Carpinteria, CA) as the primary antibody. Goat anti-rabbit conjugated to horseradish peroxidase (1:2000; catalog # 5450-0010, SeraCare, Gaithersburg, NY) was used as the secondary antibody. The immunoreaction was visualized using a 3,3-diaminobenzidine chromogen solution (DAB substrate kit; Vector Laboratories). The bright field images were captured by ScanScope CS and analyzed using ImageScope software (Aperio Technologies, Vista, CA) or ImageJ software. We recorded Iba1-positive cells area (%) and GFAP-positive astrocytes area (%) in indicated brain regions.

### Behavioral Tests

Animals were subjected to a host of behavioral tests by investigators blinded to genotype. Tests performed include the open field locomotor activity test, weight grip test, and vertical grid test to assess the motor performance of mice at days 1, 2, 3, 8, 15, and 22 after stroke. Cognitive tests were performed at baseline and day 28 post-stroke and included the novel object recognition test and Y-maze.

### Modified Vertical Grid Test

The vertical grid test is a sensitive test intended to assess neuromuscular strength and motor coordination of animals after stroke [21,22]. In previous studies, we have observed that aged animals struggle to perform this test adequately, and therefore, we modified the test to accommodate the aged animals. The vertical grid is an open frame apparatus (55 cm H × 8 cm W ×5 cm L) with a wire mesh (0.8 cm × 0.8 cm aperture) on the backside. The grid is placed in a cage filled with soft bedding material and is firmly set to an angle of 60°. Within one week prior to (baseline), and at 1, 2, 3, 8, 15, and 22 days post-pMCAO induction, each mouse was placed at the highest point on the grid facing downward and was allowed to descend the grid into the cage. A blinded investigator recorded the time required for the animal to descend. Animals were subjected to 3 trials with intervals of 30 seconds between each trial. The average of the three trials constitutes the animal’s score on the test. Animals that failed to descend the grid within 60 seconds or were unable to maintain a firm grip of the grid and fell or slid down the grid were assigned the maximum score of 60 seconds for that trial. Data for individual mice were normalized to its own baseline for analysis.

### Weight Grip Test

The weight grip test was performed with minor modifications from a previous study, as we have reported [23], to assess the muscular strength of the forepaws. Six different weights (weight 1: 16.2 g, weight 2: 30.4 g, weight 3: 44.6 g, weight 4: 58.2 g, weight 5: 71.4 g, and weight 6: 84.4 g) were prepared by attaching a metal mesh to stainless steel lines. The animals were suspended from the middle/base of the tail and allowed to grasp the first weight (weight 1). A timer starts when the mouse successfully grips the weight using its forepaws, then the animal is lifted until the steel links are completely lifted from the bench. The mouse must hold the weight for 3 seconds for the test to be successful. If the mouse was able to hold weight 1 for 3 seconds, the investigator proceeded to the next weight in sequential order. If the mouse were to drop a weight in less than 3 seconds, the test would conclude, and no further weights would be attempted. The mice were permitted three tries to successfully hold the weight for 3 seconds. A final score was tallied as a sum of the point of each weight that the mouse holds, multiplied by the number of seconds that weight was held. For example, a mouse that held up to weight 6 for 1 second is assigned a score of 1×3 + 2×3 + 3×3 + 4×3 + 5×3 + 6×1 = 51. Data for individual mice were normalized to its own baseline for analysis.

### Open Field Test

The open field test is a reliable behavioral test to assess locomotor and anxiety-like behaviors in mice. Within one week prior to stroke (baseline) and at 1, 2, 3, 8, 15, and 22 days post-stroke, the spontaneous locomotor activity of mice was measured in an open field paradigm using automated video tracking software (AnyMaze software; Stoelting, Wood Dale, IL) as previously described [24]. Mice were individually placed in an open field chamber (40 × 40 × 40 cm) with grey sidewalls and were allowed to freely explore for 10 minutes. The total distance traveled was used as indices of motor/exploratory behavior of each animal. Additionally, we recorded the amount of time spent in the center of the arena as a measure of exploratory and anxiety-like behavior. The open field arena was thoroughly cleaned with 70 % ethanol between tests. Data for individual mice were normalized to its own baseline for analysis.

### Novel Object Recognition Test

The novel object recognition (NOR) test was performed to evaluate long-term recognition memory function as described in detail in our recent studies [19,25]. Briefly, for the training session, mice were individually placed in an open field chamber (40 × 40 × 40 cm) with grey sidewalls containing two identical objects for 8 minutes and returned to their home cage. After 24 hours, the mice were exposed to the familiar arena in the presence of the original object and a novel object to test long-term recognition memory for 8 minutes. During the 8-minute test session, the time spent exploring each object was recorded. The recognition of novelty was calculated as a discrimination index: 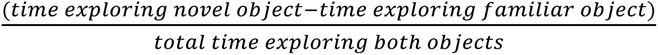. The movement of animals was videotaped by an overhead camera and analyzed using the AnyMaze software (Stoelting, Wood Dale, IL). As NOR testing was conducted at both baseline and day 28 post-stroke, two different sets of identical objects and two different novel objects were used for baseline and day 28 testing.

### Y-Maze

Spatial memory was assessed by a preference of rodents to explore novel rather than familiar places in a Y-maze. The Y-maze apparatus (15 × 3 × 5 inches; San Diego Instruments) was made of beige ABS plastic with three identical arms positioned at equal angles. In the training (“familiarization”) trial, mice were individually allowed to explore two of the three arms for a total of 5 minutes, while the third (“novel”) arm was blocked, and then returned to their home cage. After 60 minutes, the mice were placed back into the maze for 5 minutes with all three arms open. The apparatus was cleaned between trials with 70 % ethanol. Time spent in each arm was automatically determined using AnyMaze software with a camera fixed to the ceiling above the Y-maze apparatus. The number of alternations, or movements from a familiar arm to a novel arm without re-entering a familiar arm, was recorded and was used to evaluate the spatial memory performance of the experimental animals. Percent alternations is calculated by recording the % of total alternations in which the animal entered a new arm, with 50 % being due to chance.

### Statistics

The statistical analyses were performed using Prism software (GraphPad v.10). An independent unpaired Student’s t-test was performed to compare the two genotypes. Two-way ANOVA followed by Šídák’s post hoc test or repeated measures ANOVA with Tukey’s post-hoc were used for multiple comparisons. Values were expressed as mean ± SEM, and a *p*-value of less than 0.05 was considered statistically significant. The number of animals per group is clearly stated in the figure legends. Animals were assigned nondescript coded IDs upon enrollment in the study. All studies were performed by investigators blinded to the treatment of the animals in analyses.

## Results

### *Ripk2^-/-^* aged mice have reduced infarct volume compared to aged wildtype (WT) controls

To assess the effects of *Ripk2* genetic deficiency on infarct volume in aged mice, we performed permanent middle cerebral artery occlusion (pMCAO) in aged wildtype (WT) and *Ripk2^-/-^* mice. **Figure 1A** shows representative cresyl violet stained sections from both WT and *Ripk2^-/-^* mice at 28 days following pMCAO. Infarct volume is quantified in **Figure 1B**, and *Ripk2^-/-^*aged mice displayed a significantly reduced infarct volume compared to WT controls.

**Figure 1:**
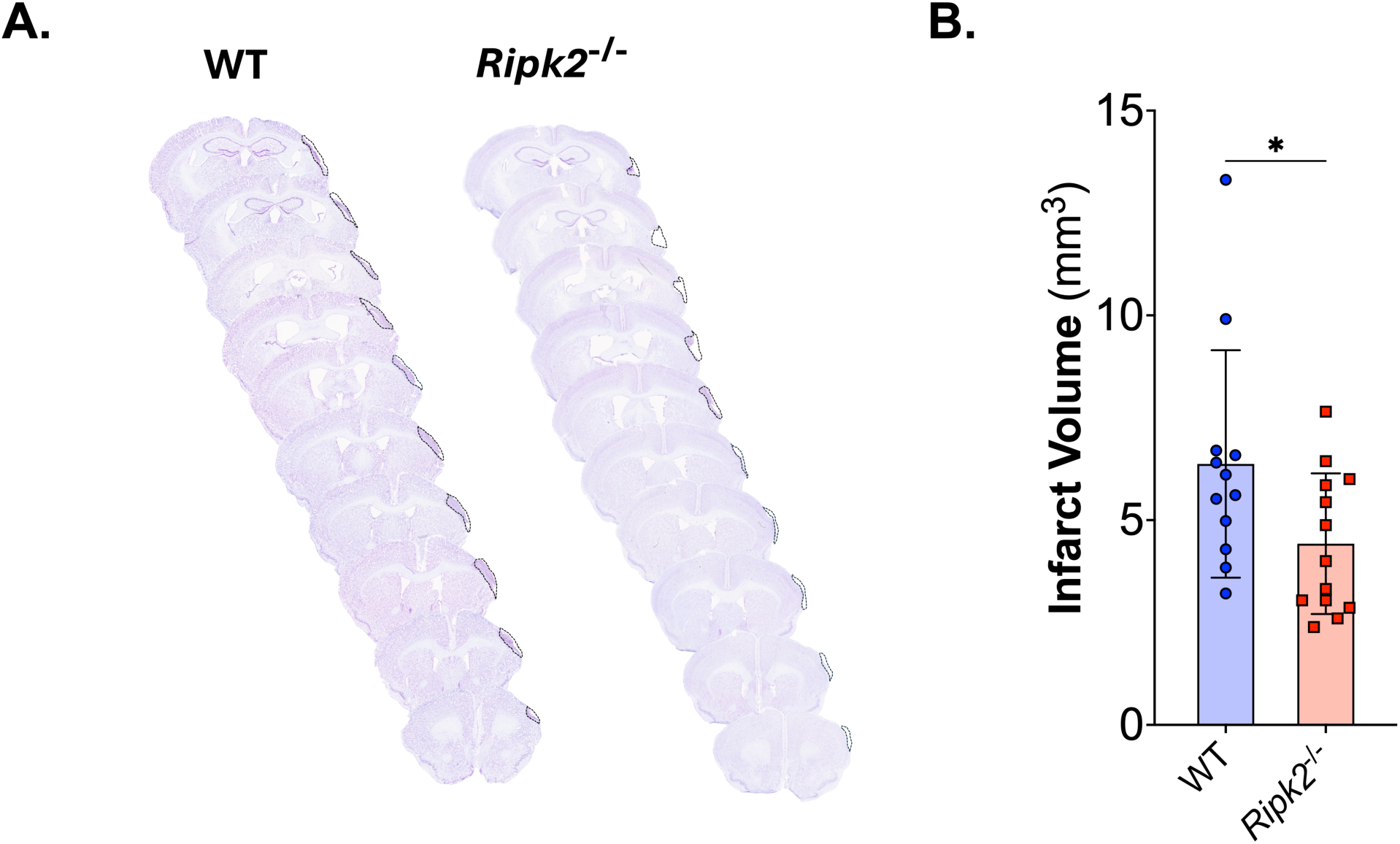
*Ripk2^-/-^*mice have reduced infarct volume at 28 days than aged WT mice. **A)** Representative cresyl violet stained sections of brains from either wildtype (WT) or *Ripk2* deficient (*Ripk2^-/-^*) mice. Sections cut at 30 µm using a cryostat, stained with cresyl violet, and imaged using an Aperio slide scanner. **B)** Quantification of infarct volume shows significant reduction in infarct volume in the aged *Ripk2^-/-^*mice compared to WT mice. *n* = 12-13/group. Differences determined by two-tailed, unpaired *t*-test; *t*(23) = 2.128, *p* = 0.0443.

### *Ripk2^-/-^* mice show improved performance in motor behavior tests following pMCAO

For 28 days following pMCAO, mice were subjected to longitudinal locomotor behavior tests, including the vertical grid, weight grip, and open field tests. Tests were also performed at baseline prior to pMCAO for training and to determine if there were baseline differences between the *Ripk2^-/-^*mice and WT controls. At baseline, aged *Ripk2^-/-^* mice took more time to descend the vertical grid than WT aged controls (**Figure 2A**). Additionally, at baseline, aged *Ripk2^-/-^* mice were able to hold less weight in the weight grip test and traveled less distance in the open field test than aged WT controls (**Figures 2C and 2E**). Because of these baseline differences between genotypes, longitudinal data were normalized to the baseline data. When normalized to baseline data, aged *Ripk2^-/-^*mice were able to descend the vertical grid more quickly (**Figure 2B**) and hold more weight (**Figure 2D**) than aged WT controls after stroke. There were no differences between groups in the longitudinal open field test total distance traveled (**Figure 2F**); however, *Ripk2^-/-^* mice spent less time in the center of the open field arena than WT controls at baseline and days 1 and 3 post-stroke (**Supplementary Figure 1)**.

**Figure 2:**
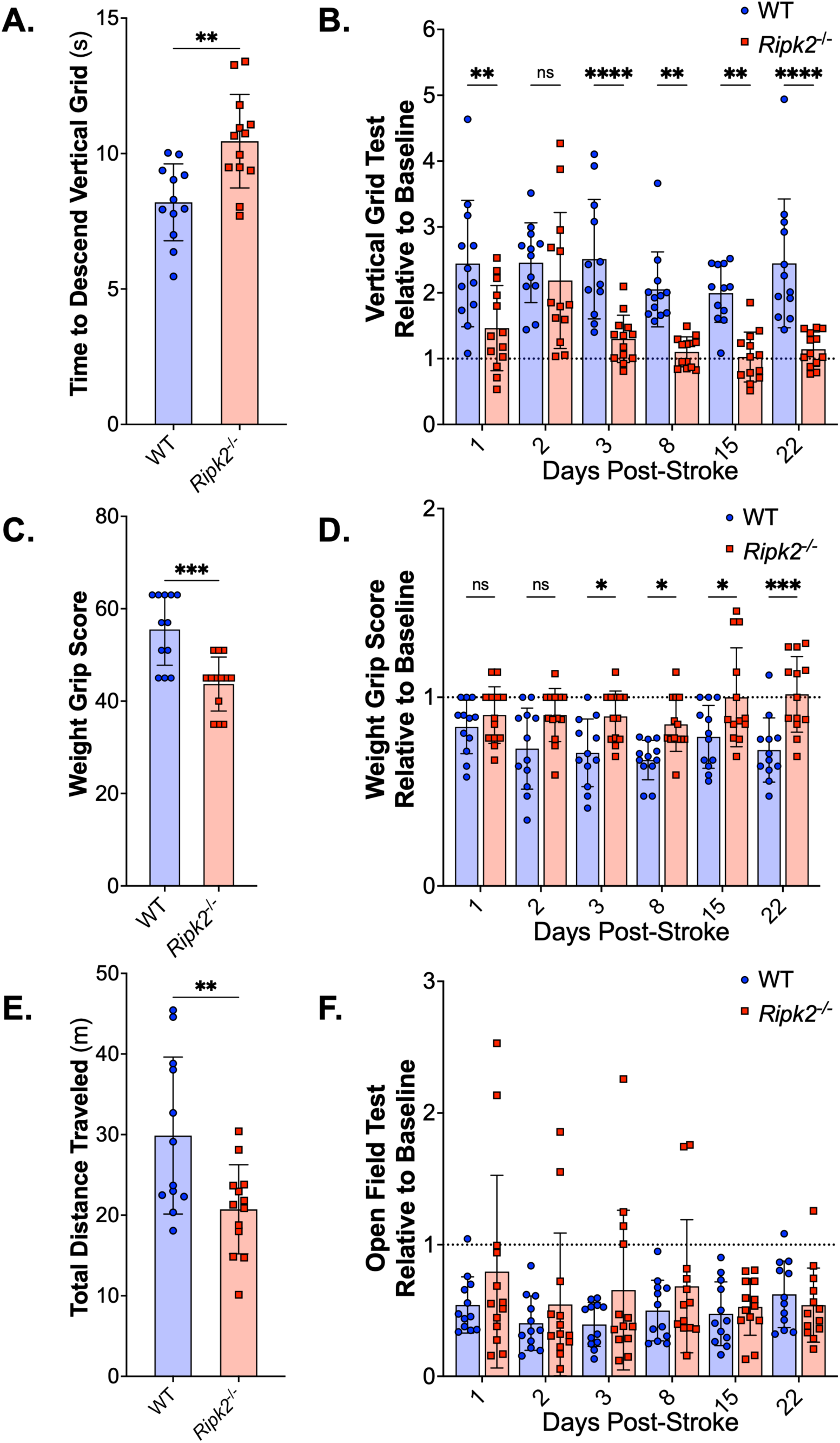
*Ripk2^-/-^*aged mice show improved function on behavior tests compared to aged WT controls. **A)** Time to descend the vertical grid at baseline was significantly different between groups, with *Ripk2^-/-^* mice taking more time to descend the grid. **B)** When normalized to baseline data for each genotype, *Ripk2^-/-^* mice show less deficit in descending the vertical grid than WT controls at days 1, 3, 8, 15, and 22. **C)** *Ripk2^-/-^* mice had lower weight grip scores than WT controls at baseline. **D)** *Ripk2^-/-^* mice have higher weight grip scores normalized to baseline than WT controls at days 3, 8, 15, and 22 following pMCAO. **E)** *Ripk2^-/-^* mice traveled less distance than WT controls at baseline in the open field test. **F)** No difference between genotypes in baseline normalized total distance traveled in the open field test following pMCAO. *n* = 12-13/group. Differences determined by two-tailed, unpaired *t*-test (A, C, E). Differences determined by two-way, repeated measures ANOVA with Tukey’s post-hoc (B, D, F). * = *p* < 0.05, ** = *p* < 0.01, ***= *p* < 0.001, **** = *p* < 0.0001.

### *Ripk2^-/-^* mice perform poorly in cognitive assays before and after pMCAO

Because we were assessing mice of different genotypes at advanced age in a longitudinal study with cognitive tests, it was important to assess cognitive abilities of the mice prior to the induction of the pMCAO model. A baseline novel object recognition (NOR) test was performed by exposing mice to two identical objects during the training phase (**Figure 3B**) before swapping one of the objects for a novel object in the testing phase (**Figure 3C**, schematic shown in **Figure 3A**). Generally, mice should spend more time with the novel object in the testing phase as an indication that they remember the old object and recognize the new object. However, in the baseline testing phase (**Figure 3C**), *Ripk2^-/-^* mice did not show increased exploratory time with the novel object, but the aged WT control mice spent significantly more time with the novel object. This is further illustrated in the discrimination index (**Figure 3D**), which shows higher object discrimination in the aged WT controls than in the aged *Ripk2^-/-^*mice at baseline. Twenty-eight days following pMCAO, the NOR test was repeated with entirely new objects. During the training phase (**Figure 3E**), there were no differences between genotypes in time spent with objects. During the day 28 testing phase (**Figure 3F**), the *Ripk2^-/-^* mice spent less time with the novel object than with the familiar object, while the aged WT control mice showed no difference between the objects. There was no difference in discrimination indices between the genotypes (**Figure 3G**), indicating that both aged WT and aged *Ripk2^-/-^* mice had cognitive deficits following pMCAO.

**Figure 3:**
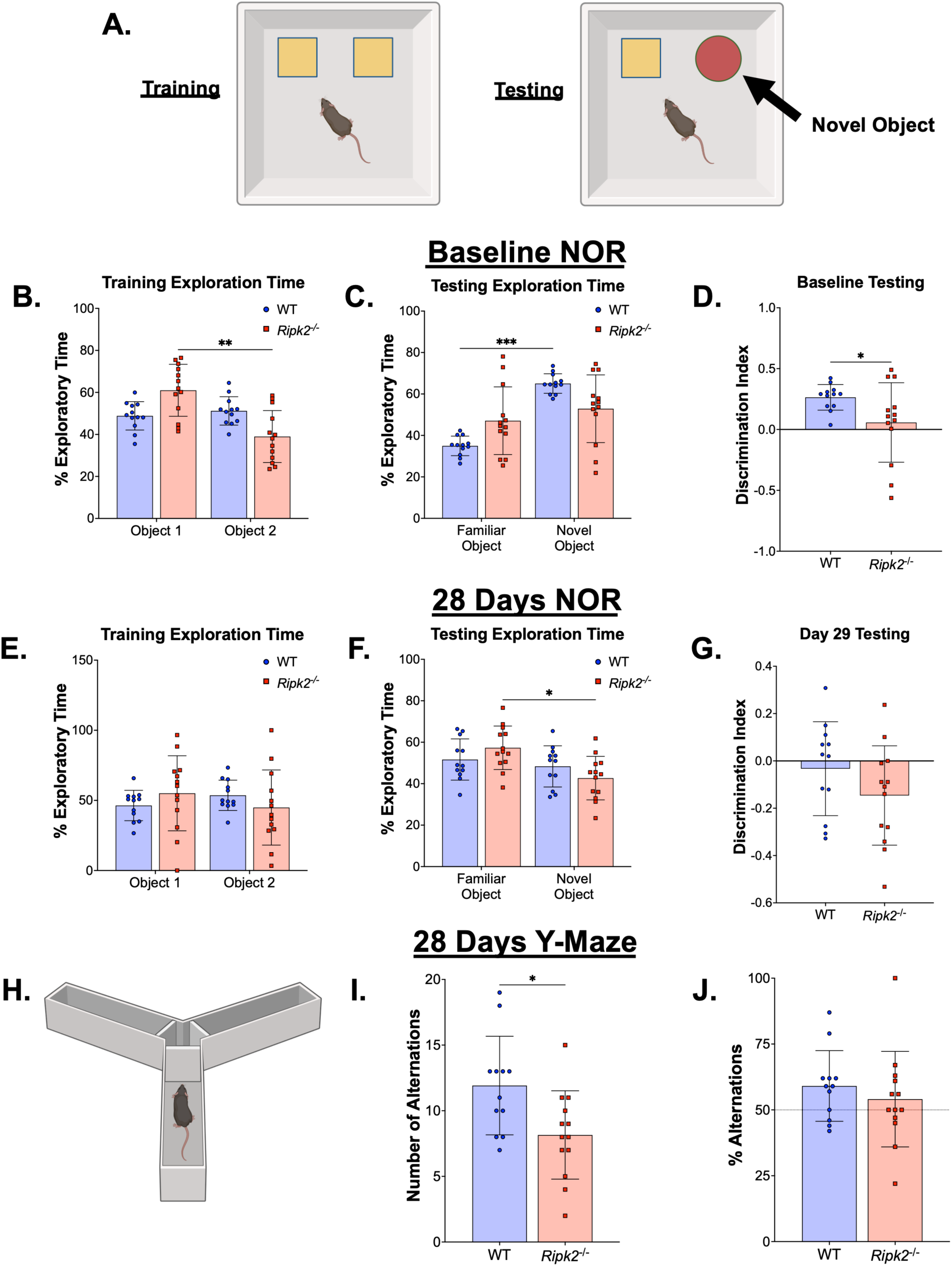
Cognitive testing reveals *Ripk2^-/-^* mice display cognitive deficits before stroke and do not show worsened cognitive deficits post-stroke. **A)** Schematic depiction of the novel object recognition (NOR) test during the training (left) and testing (right) phases. **B-D)** Baseline NOR training (B), testing (C), and discrimination index (D) conducted in naïve animals. **E-G)** NOR testing with the training phase (E) conducted on day 27 and testing (F) on day 28, resulting in a discrimination index (G). **H-J)** Y maze cognitive test performed 28 d post-stroke. **H)** Schematic of the Y maze test paradigm. **I)** The number of alterations per animal. **J)** The percent alterations per animal. n=12-13/group. (B-C, E-F) Differences were determined by two-way ANOVA with Šídák’s post-hoc. (D, G, I-J) Differences were determined by Student’s t-test. * *p* < 0.05, ** *p* < 0.01, *** *p* < 0.01, **** *p* < 0.0001.

Additionally, on day 28 we performed the Y-maze test (**Figure 3H**) to assess cognitive function following pMCAO. Mice were trained for 5 minutes with two arms of the maze open, and one hour later mice were placed back into the apparatus with all three arms open. Aged *Ripk2^-/-^* mice showed fewer total alternations than aged WT mice (**Figure 3I**); however, there were no differences in % alternations between the two genotypes, with both performing slightly higher than chance (**Figure 3J**).

### Aged *Ripk2^-/-^* mice show no difference in GFAP expression and less Iba1 reactivity than aged WT controls following pMCAO

To assess gliosis following pMCAO in aged WT and aged *Ripk2^-/-^*mice, brain sections were stained for Iba1, a marker for microglia/macrophages, and glial fibrillary acidic protein (GFAP), a marker for astrocytes. Whole brain images of Iba1 expression are shown in **Figure 4A**, with rectangles indicating the areas of cortex and subcortex quantification (**Figures 4B** and **4D**). Staining was quantified using ImageJ and is shown in **Figures 4C** and **4E**. Aged WT control animals showed increased Iba1 expression in the ipsilateral cortex compared to the contralateral cortex following pMCAO, while aged *Ripk2^-/-^* mice do not. Additionally, aged WT mice have increased expression of Iba1 in the ipsilateral cortex compared to aged *Ripk2^-/-^* mice, potentially showing a negative regulation role of RIPK2 in microglial/macrophage activation following stroke.

**Figure 4:**
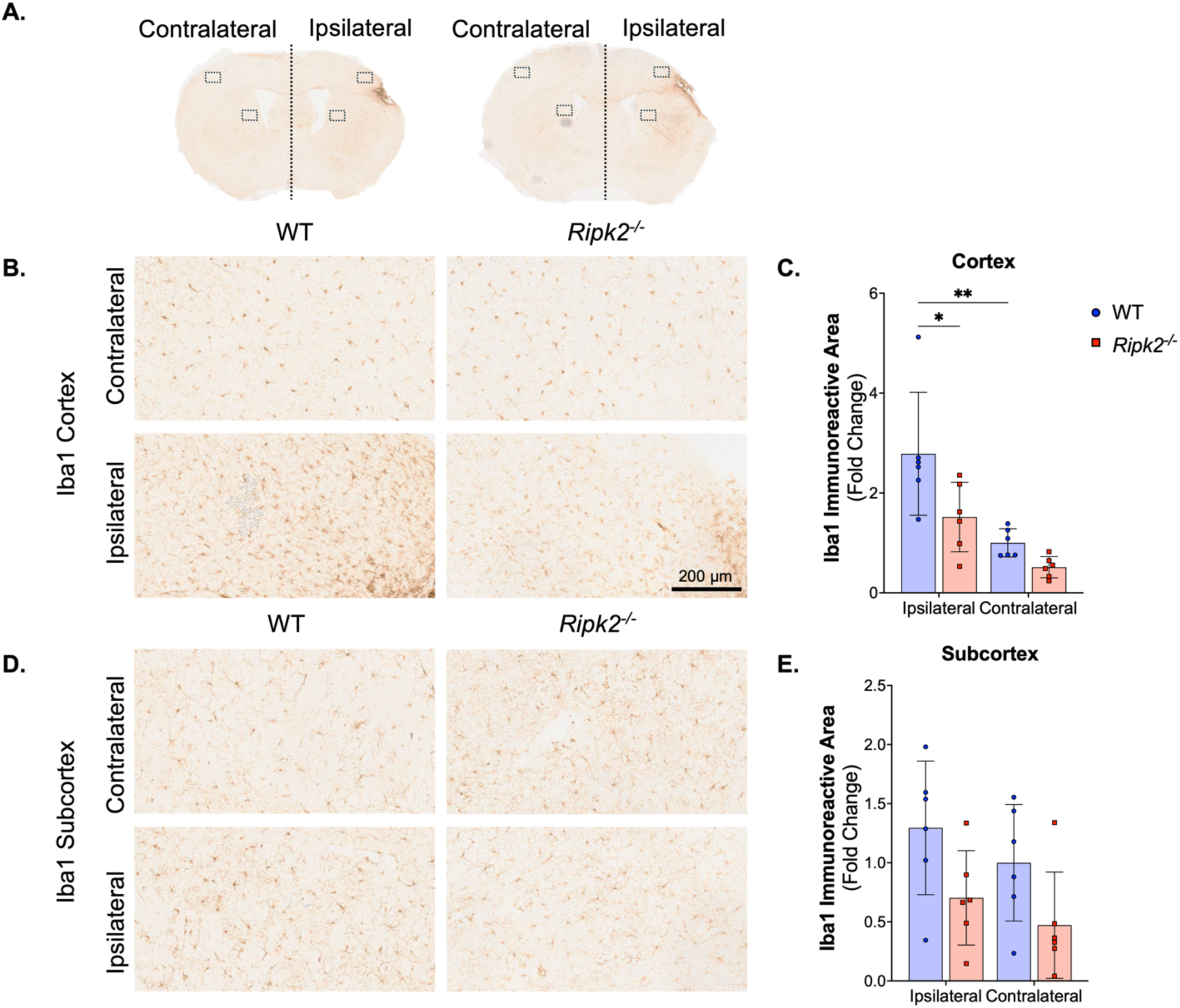
*Ripk2^-/-^*mice have less Iba1 staining than WT control mice in the ipsilateral cortex following pMCAO. **A)** Representative 30 µm sections show Iba1 staining following pMCAO in WT (left) and *Ripk2^-/-^* mice (right). **B)** Images taken from the contralateral (top) and ipsilateral (bottom) cortex. Scale bar is 200 µm. **C)** Quantification of Iba1 immunostaining shown in panel B. WT mice show increased Iba1 staining in the ipsilateral cortex compared to the contralateral cortex, and *Ripk2^-/-^* mice show decreased Iba1 staining in the ipsilateral cortex compared to WT controls. **D)** Images taken from the contralateral (top) and ipsilateral (bottom) subcortex. **E)** Quantification of Iba1 immunostaining shown in panel D shows no differences between genotypes or hemispheres in Iba1 immunostaining in the subcortex. *n* = 6/group. Differences determined by two-way ANOVA with Tukey’s post-hoc. * *p* < 0.05, ** *p* < 0.01.

Whole sections were imaged using an Aperio slide scanner and images of contralateral and ipsilateral cortex and subcortex were quantified using ImageJ. Whole brain images of GFAP expression are shown in **Figure 5A**, with rectangles indicating the areas of cortex and subcortex quantification (**Figures 5B** and **5D**). Staining was quantified using ImageJ and is shown in **Figures 5C** and **5E**. Both aged WT and aged *Ripk2^-/-^* mice had increased GFAP expression in the ipsilateral cortex compared to the contralateral cortex following pMCAO, with no differences between genotypes. There were no differences between genotypes or hemispheres with GFAP expression in the subcortex.

**Figure 5:**
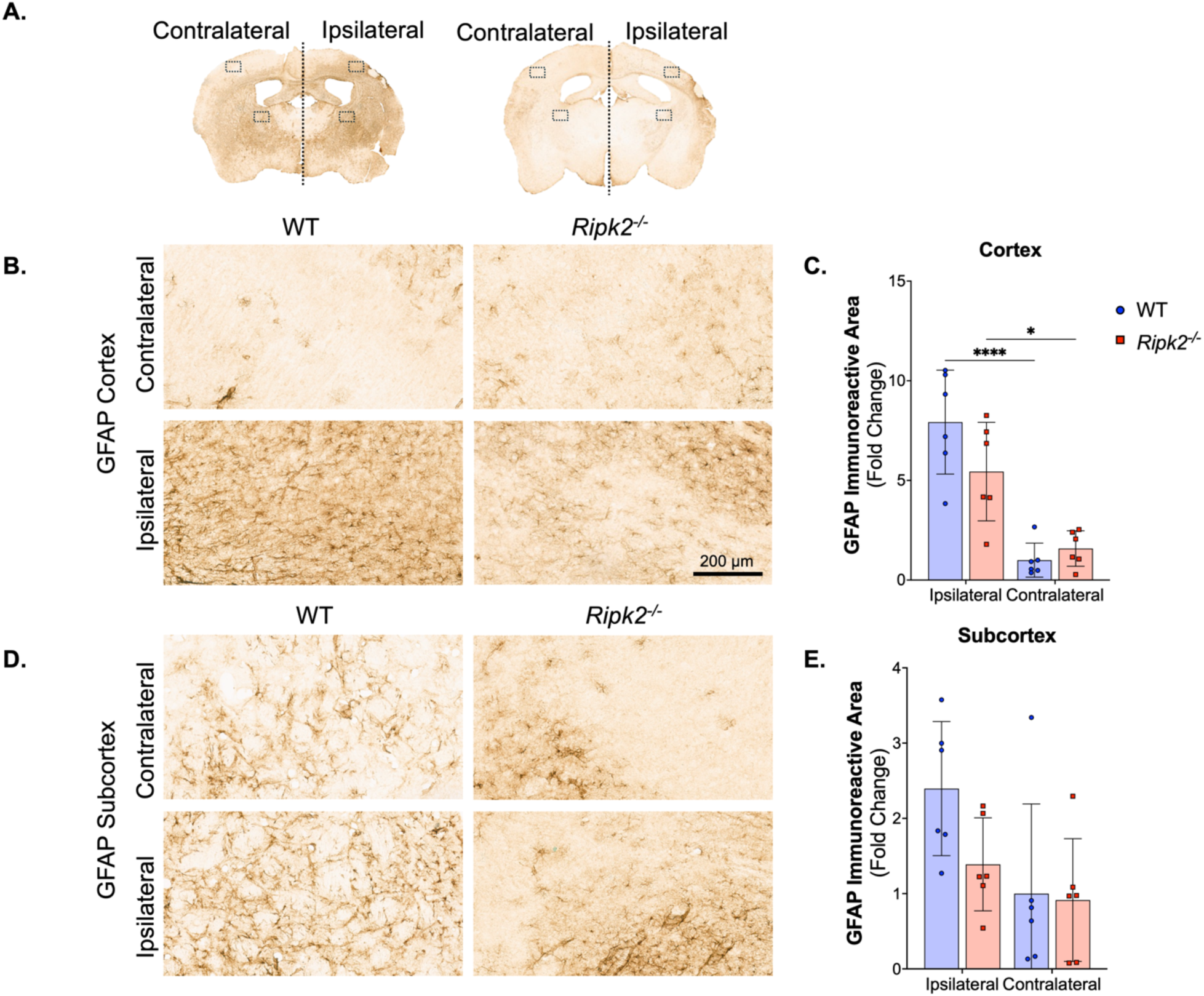
Both WT and *Ripk2^-/-^*mice show increased GFAP staining in the ipsilateral cortex with no difference between genotypes. **A)** Representative 30 µm sections show GFAP staining following pMCAO in WT (left) and *Ripk2^-/-^* mice (right). **B)** Images taken from the contralateral (top) and ipsilateral (bottom) cortex. Scale bar is 200 µm. **C)** Quantification of GFAP immunostaining shown in panel B. Both WT and *Ripk2^-/-^* mice show increased GFAP staining in the ipsilateral cortex compared to the contralateral cortex. **D)** Images taken from the contralateral (top) and ipsilateral (bottom) subcortex. **E)** Quantification of GFAP immunostaining shown in panel D shows no differences between genotypes or hemispheres in GFAP immunostaining in the subcortex. *n* = 6/group. Differences determined by two-way ANOVA with Tukey’s post-hoc. * *p* < 0.05, **** *p* < 0.0001.

## Discussion

In the current study, we utilized a genetic approach to assess the role of RIPK2 in neuroinflammation following the pMCAO model of stroke in aged mice. Our results show that RIPK2 plays an important role in the post-stroke pathology that worsens infarct size and behavioral outcomes. This is illustrated by our data showing that aged *Ripk2^-/-^* mice had smaller infarct volumes than aged WT control mice. Aged *Ripk2^-/-^* mice also descended the vertical grid more quickly and held more weight following stroke than the WT controls. Lastly, *Ripk2^-/-^* mice had less Iba1 staining in the ipsilateral cortex than the aged WT control mice, suggesting less microgliosis and/or infiltration of peripheral monocytes/macrophages. There was also a strong trend towards decreased GFAP staining in both the ipsilateral cortex and subcortex of *Ripk2^-/-^* mice compared to WT controls, but it did not reach significance.

In our cognitive tests, aged *Ripk2^-/-^* mice performed worse than WT controls both at baseline and following pMCAO. One possible explanation for this is that aged *Ripk2^-/-^* mice may have higher anxiety-like behavior than WT controls. This is further illustrated by our open field test data, as the *Ripk2^-/-^* mice spent less time in the center of the arena than the WT controls both at baseline and following stroke. Anxiety-like behavior in rodents can present in a variety of ways, but in tests like the Y-maze and NOR tests, it can present as neophobia or avoidance of novel stimuli [26,27], while in the open field test, it presents as avoidance of the center of the arena. To our knowledge, *Ripk2^-/-^* mice have not directly been assessed for anxiety-like behavior, but a previous report using young *Ripk2^-/-^*mice reported that their anxiety behavior was normal [28]. The increased anxiety seen in our data could, therefore, be a result of the interplay between the lack of functional RIPK2 and advanced age in the *Ripk2^-/-^* mice used in this study. Additionally, previous studies have shown that RIPK2 regulates neuronal survival during development through its interactions with the p75 neurotrophin receptor, which functions to regulate cell death, in cerebellar granule neurons and Schwann cells [29,30]. In another study using NOD1/NOD2 double knockout mice, it was reported that the NOD double knockout mice had higher anxiety-like behavior and worsened performance on cognitive tests like novel object recognition after stress [31], further implicating the role of NOD-signaling and RIPK2 in neuronal survival, anxiety-like behavior, and cognitive impairment. Future studies should investigate the precise role of RIPK2 and NOD signaling in post-stroke cognitive impairment.

Previous studies in our lab have explored the role of RIPK2 in post-stroke neuroinflammation mechanistically by using genetic models and pharmacological inhibitors of RIPK2 in young animals [15,16]. We have shown that *Ripk2^-/-^* global knockout mice and *Ripk2^-/-^*microglial conditional knockout mice have smaller infarct volumes and improved behavioral outcomes following transient MCAO, with a greater effect seen in the global knockout mice [16]. We have also shown in young mice that pharmacologically inhibiting RIPK2 following stroke is neuroprotective, reduces infarct volume, and improves behavioral outcomes in acute studies [15]. Because stroke predominantly occurs in aged patients [1], it was important to validate our previous findings in aged mice. Additionally, we opted to do a longitudinal study in the aged *Ripk2^-/-^* mice to assess long-term recovery and cognitive function following stroke. This is incredibly important due to the increased prevalence of post-stroke cognitive impairment. Some studies estimate that as many as ∼ 60% of stroke survivors experience cognitive impairments [32–35]. We also decided to use the clinically relevant pMCAO model of ischemic stroke because many large vessel occlusions are not fully resolved in the clinical stroke population. The current study expands our previous work by exploring the role of RIPK2 in aged mice for 28 days after pMCAO.

Inflammatory signaling plays an important role in the ischemic cascade. It is well documented that inhibiting and modulating either neuroinflammation or systemic inflammation can improve outcomes in rodent models of ischemic stroke. Specific mediators of the inflammatory cascade have been targeted in preclinical research as potential drug targets, such as the NF-κB inflammatory pathway [8,36,37]. RIPK2 is involved in inflammatory signaling upstream of both NF-κB and MAPK, two major pathways promoting inflammatory gene transcription, making it an attractive candidate for reducing inflammation [8,9,11]. However, it is also known that chronic inhibition of inflammation can have negative effects, as inflammatory signaling promotes repair in the later phase of stroke recovery.

A limitation of this study is that we used only aged male mice. We have previously shown that aged female *Ripk2^-/-^* mice have smaller infarct volumes in acute studies [16], but future studies will need to assess the effects of *Ripk2^-/-^* longitudinally in aged mice of both biological sexes, since women have been shown to have worsened outcomes and increased mortality following stroke [38]. Additionally, future studies should use pharmacological inhibitors of RIPK2 long-term following stroke to further investigate the therapeutic potential of RIPK2 inhibition to reduce inflammation and improve outcomes following ischemic stroke.

## Conclusions

This study utilized aged *Ripk2^-/-^* mice to assess the role of RIPK2 in post-stroke neuroinflammation at advanced age. We utilized the pMCAO model and examined the effects of *Ripk2^-/-^* on behavior outcomes, infarct volume, and microglial/macrophage and astrocytic activation at 28 days post-stroke. This study is significant because it uses aged animals, an important consideration for experimental stroke research, to investigate the effects of RIPK2 signaling on post-stroke inflammation and its effects on infarct volume and behavioral outcomes.

## Acknowledgments

This work was supported by NIH grants R01NS109816 and R01NS129136 from the NINDS to ECJ, a Transformational Project Award from the American Heart Association to ECJ (award # 971058), and a Predoctoral Fellowship from the American Heart Association to JL (award # 915693). This work is also supported by McKnight Brain Institute Gator NeuroScholars funding to JAH and a Postdoctoral Fellowship from the American Heart Association to JAH (25POST1375222).

## Authors Contributions

Conceptualization: JAH, JL, ECJ. Methodology: JAH, JL, REG, SMS, LL, CY, ECJ. Software: JAH, JL. Formal analysis: JAH, JL, ECJ. Data curation: JAH, JL, ECJ. Visualization: JAH, JL, ECJ. Writing – original draft: JAH, JL, ECJ. Writing – review and editing: All authors provided input. Supervision: ECJ. Funding acquisition: JAH, ECJ. All authors read and approved the final manuscript.

## Data availability

The original data supporting this study’s findings are available from the corresponding author upon reasonable request.

---

**Supplementary Figure 1:**
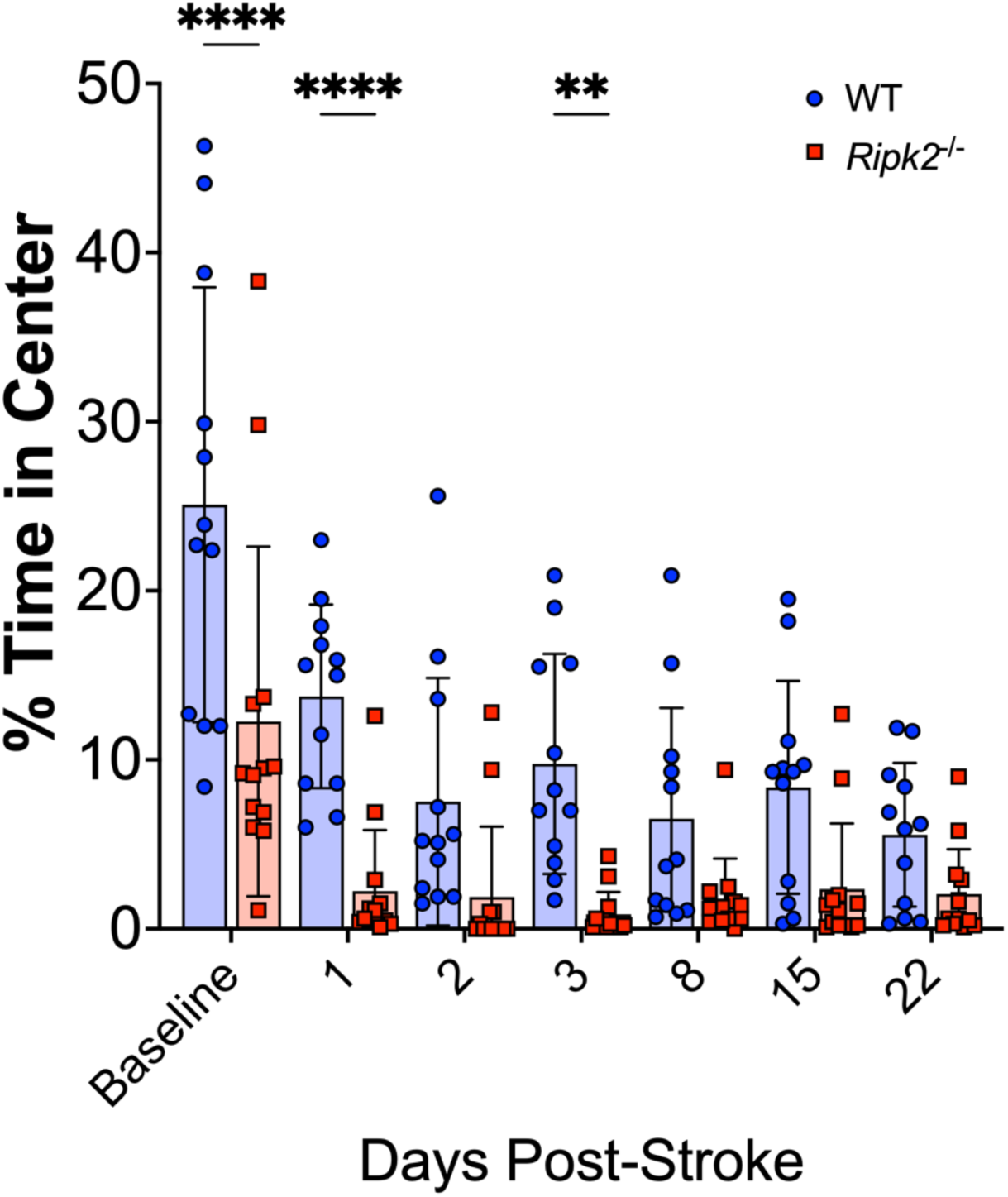
Ripk2-/- mice spend less time in the center of the open field chamber than aged WT control mice. Quantified percent time in center of the open field arena shows decreased time in center for *Ripk2^-/-^* mice compared to aged WT control mice. Differences detected using two-way repeated measures ANOVA with Šídák’s post-hoc. ** *p* < 0.01, **** *p* < 0.0001. At baseline, before permanent middle cerebral artery occlusion induction, *Ripk2^-/-^*mice spent less time in the center of the open field arena than aged WT control mice. This also happened at days 1 and 3 post-stroke, with strong trends throughout the study. This could indicate higher anxiety-like behavior in *Ripk2*^-/-^ mice than in the WT control mice.

